# A dual role for CTCF in development

**DOI:** 10.64898/2026.03.17.712290

**Authors:** Noemi Alonso Saiz, Moreno Martinovic, Miguel Rubio, Pinak Samal, Stefan Giselbrecht, Luca Braccioli, Elzo de Wit

## Abstract

CTCF is an essential DNA binding protein whose absence leads to embryonic lethality. CTCF is primarily known for its role in 3D genome organization where its N-terminal domain interacts with cohesin to anchor chromatin loops. How CTCF facilitates proper embryonic development remains unclear, necessitating temporal control to resolve its stage-specific functions. By combining gastruloids, an *in vitro* model of embryonic development, with a degron system to rapidly deplete CTCF at defined timepoints, we show that early CTCF depletion impairs early gastruloid morphogenesis. Surprisingly, ATAC-seq and time-resolved RNA-seq revealed that differentiation was unaffected. CTCF binding is strongly enriched at promoters of downregulated genes. Re-expression of a CTCF variant with an N-terminal truncation, incapable of looping, was sufficient to rescue the expression of CTCF-promoter bound genes and the defects in morphogenesis. However, extended culture (up to 168 hours) of gastruloids reconstituted with N-terminal truncated CTCF led to their collapse. Our work shows that CTCF has a dual function in early mammalian development: at early stages CTCF regulates developmentally important genes through promoter binding, while at later stages its looping function is required for correct development.

**Highlights:**

- CTCF is essential for gastruloid morphogenesis but dispensable for cell differentiation
- CTCF activates genes through promoter binding
- CTCF promoter target regulation drives *in vitro* gastrulation
- Post-gastrulation development *in vitro* is driven by CTCF’s looping function

## Introduction

During the first stages of embryonic development, the totipotent zygote undergoes successive rounds of cell division and differentiation to form the blastocyst. Before implantation, the blastocyst is composed of the trophectoderm and the inner cell mass. The latter will give rise to the pluripotent epiblast^1^. Following implantation, the epiblast undergoes gastrulation, during which the primitive streak emerges, and the three germ layers — ectoderm, mesoderm and endoderm — are established. This culminates in the formation of the basic body plan of the embryo^2,3^.

The development of mammalian embryos in the uterus complicates the study of early development *in vivo*. In recent years, gastruloids have emerged as an *in vitro* model for gastrulation and early organogenesis^4–17^. Mouse gastruloids develop from aggregates of embryonic stem cells that self-aggregate and produce cell types and gene expression patterns that resemble the ones found during early embryonic development. Initially, gastruloids grow in differentiation media. At 48h, they are stimulated with the Wnt-agonist Chiron, inducing symmetry breaking, polarization and elongation. After 24h, Chiron is removed and gastruloids are maintained in differentiation media. While continuing their differentiation trajectory, they display key morphogenetic pathways similar to those driving the development of the primitive streak in the mouse embryo^2,4,12,13,15^.

Expression of key developmental regulators depends on the correct organization of the genome^18–26^. The zygotic genome lacks organizational features commonly found in developed cell types^27–30^. However, following the first cell divisions, topologically associating domains (TADs) and DNA loops are established^27–30^. TADs are formed through the loop extrusion mechanism, whereby a cohesin complex extrudes DNA until it encounters a bound CTCF site. Extrusion is stalled upon interaction of cohesin with the N-terminal domain of CTCF. In this way, CTCF can act as a facilitator or insulator of enhancer-promoter communication^31–49^, and contribute to precise developmental gene regulation^23,26,50,51^.

The role of CTCF as an insulator is important for several developmental processes. The removal or alteration of a single or multiple CTCF boundaries can lead to developmental disorders. In neurons, CTCF binds to the protocadherin (Pcdh) cluster, allowing the stochastic expression of each Pcdh gene to promote axon self-avoidance^22^. During mouse embryonic development, the rearrangement of the TAD containing the *Epha4* locus results in limb malformations^51^. Moreover, during early embryonic development, CTCF controls the temporal and sequential activation of the Hox genes, which are essential for the establishment of the rostrocaudal axis^23,26^. Furthermore, global alterations in CTCF function during development have severe consequences for the organism. In humans, mutations in the CTCF coding sequence result in neurodevelopmental disorders^52–55^. In mouse and zebrafish, zygotic CTCF knockouts are embryonically lethal^56,57^. Both embryos and cell lines deficient for CTCF present a nearly complete loss of TADs. Despite its essential role at both the cellular and the organismal level, the direct role of CTCF in regulating gene expression seems limited^39,58^.

Because CTCF is essential and ubiquitously expressed, a system that enables temporal control of CTCF levels is required to disentangle its roles at distinct developmental stages. Here, we use gastruloids grown from mouse embryonic stem cells (mESCs) engineered with a degron system to selectively deplete CTCF at specific times of gastruloid development..

We find that while CTCF is required for morphogenesis, it is not essential for cell differentiation. Through rescue experiments we show that CTCF promoter binding is required for the early stages of gastruloid development. This function is independent of its looping role. However, during later stages of gastruloid development the looping function of CTCF is required for proper development. Our experiments in gastruloids demonstrate a dual role for CTCF regulating different stages of early embryonic development.

## Results

### CTCF is required for gastruloid morphogenesis but not for cell differentiation

To study the role of CTCF in early development, we combined two powerful technologies: mouse gastruloids and degrons^47,59–62^. We grew gastruloids by aggregating 400 mESCs where CTCF has been endogenously tagged with an auxin-inducible degron (AID)^39,59^ (CTCF-AID, Figure 1A). This approach allows selective depletion of CTCF at different timepoints during gastruloid development by addition of auxin (IAA) in the medium (Figure 1A). We confirmed that CTCF was rapidly degraded in mESCs, as the protein was undetectable after 1 hour of IAA treatment (Figure S1A). As control, we used mESCs with the same genetic background but where CTCF is not tagged (parental, PT). Gastruloids successfully developed from both CTCF-AID and PT cells (Figure 1B). Without treatment (not-treated, NT), CTCF-AID and PT gastruloids were similar in size and showed proper elongation (Figure 1B-C), demonstrating that the CTCF degron cells can form gastruloids. To determine how gastruloids develop in the absence of CTCF, we treated gastruloids with IAA either from 48 or 72 hours of gastruloid development (Figure 1B). Gastruloids depleted of CTCF from 48 hours onwards, failed to elongate and yielded significantly smaller gastruloids (Figure 1B-C), consistent with the essential role of CTCF in early development. Surprisingly, depleting CTCF only 24 hours later (+IAA@72h) had a minimal effect on gastruloid elongation (Figure 1B-C). These results show that CTCF is essential for *in vitro* gastrulation during the early stages of gastruloid development, recapitulating the *in vivo* findings in CTCF KO embryos^29,57,63^.

**Figure 1.**
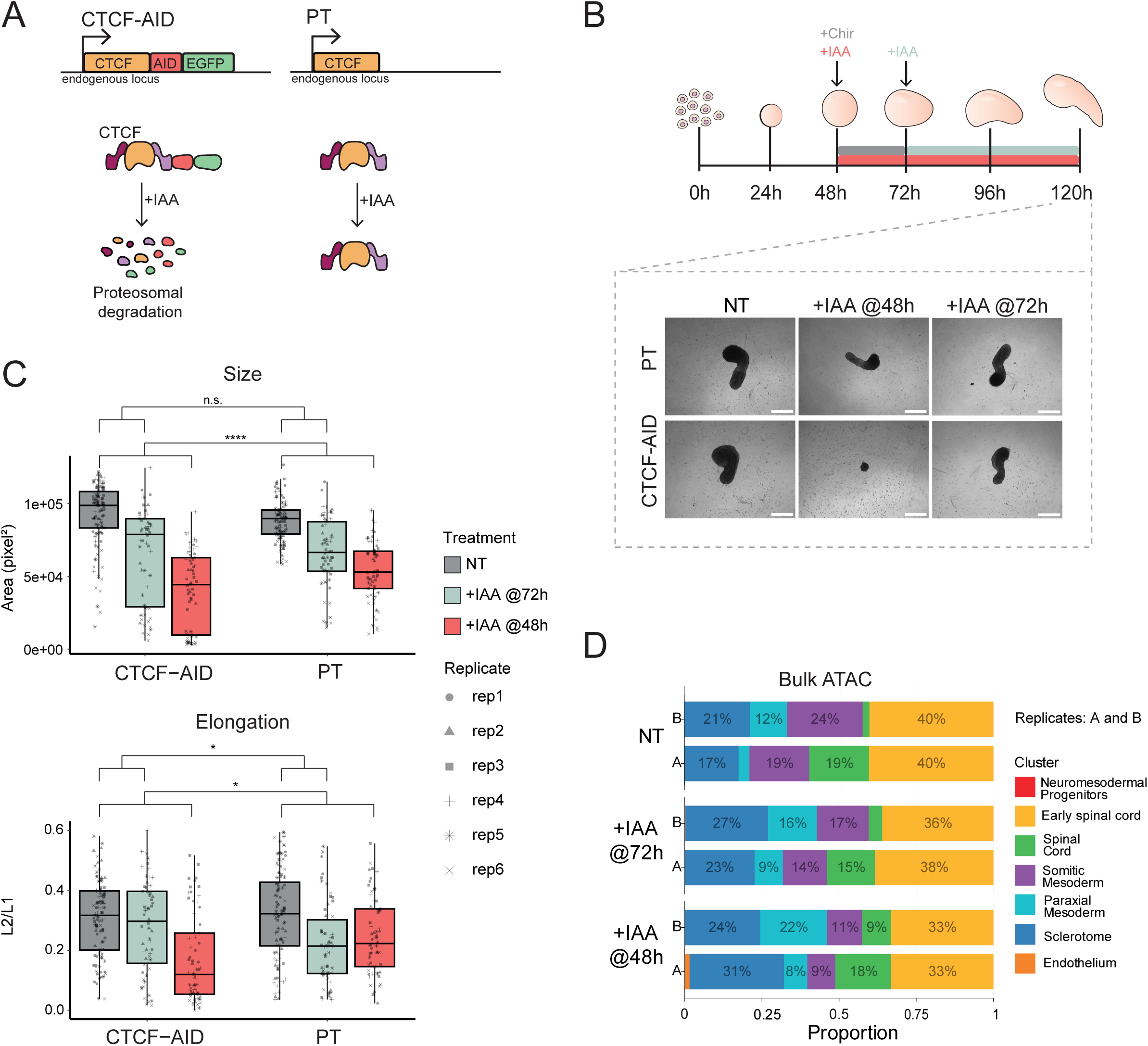
CTCF is required for development but not for differentiation A. Schematic representation of the endogenous CTCF locus or the CTCF protein in the two cell lines used in this study: CTCF-AID and parental (PT). B. Gastruloid culture timeline. +Chir and +IAA indicate treatment with CHIR99021 and IAA respectively. Bright field images of representative 120 hours gastruloids were taken using a wide field microscope. Scale bar: 300um. C. Size and elongation quantification of 120h gastruloids. Size was calculated in pixel^2^. Elongation is represented by the ratio between the L2 and L1 LOCO-EFA coefficients^95^. Values were obtained using MOrgAna^94^. Difference in treatment effects was tested using linear models with Benjamini-Hochberg multiple testing correction (n.s. = p > 0.05; * = p ≤ 0.05; ** = p ≤ 0.01; *** = p ≤ 0.001; **** = p ≤ 0.0001). D. Bulk ATAC deconvolution on 120 hours gastruloids with two independent replicates per treatment.

The developmentally relevant CTCF depletion timepoint (48 hours) coincides with the activation of Wnt signalling by Chiron. Wnt signaling induces the expression of Brachyury (T or TBXT) in a subset of cells that drive gastruloid polarization between 48 and 72 hours^64^. Over this period, the gastruloid transitions from a largely pluripotent population of SOX2-positive cells at 60 hours to a more heterogeneous mixture of primed cells displaying variable SOX2 expression by 72 hours. We investigated whether CTCF depletion from 48 hours affects the onset of these expression changes and gastruloid polarization. We performed SOX2 and T immunofluorescence staining on CTCF-AID gastruloids treated with IAA at 48 hours and collected gastruloids at 60 hours and 72 hours (Figure S1B). CTCF-AID-treated gastruloids showed the same SOX2 and T expression pattern as the untreated (NT) controls, demonstrating that the onset of symmetry breaking and the first cell specification events can occur in the absence of CTCF.

Given the lack of gastruloid elongation we wondered whether CTCF depletion from 48 hours can impair differentiation events or alter cell-type distribution in gastruloids. To answer this, we determined the chromatin accessibility landscape by performing Assay for Transposase Accessible Chromatin followed by high-throughput sequencing (ATAC-seq) on 120 hours gastruloids. We used our previously generated single-cell ATAC-seq data to deconvolve the bulk signal^5,65^. In this manner, we determined the cell lineages present in 120 hours gastruloids and their relative abundance (Figure 1D). CTCF depletion from 72 hours had minimal effect on cell type composition compared to the NT controls (Figure 1D), consistent with their morphological similarity (Figure 1B). Surprisingly, gastruloids depleted of CTCF from 48 hours, which showed severe elongation defects (Figure 1B), had a similar distribution of the inferred cell types (Figure 1D). Taken together, these results show that CTCF is not required for cell differentiation in gastruloids and that we can uncouple differentiation from morphogenesis.

### Effects of CTCF depletion on the temporal regulation of developmental pathways

CTCF is essential for morphogenesis between 48 and 72 hours of gastruloid development. The severe morphological defects are likely a consequence of changes in gene expression upon CTCF depletion. To better understand which genes are regulated by CTCF in gastruloids, we performed a high-resolution time-resolved bulk RNA-seq experiment. We cultured gastruloids derived from PT and CTCF-AID cells, with or without IAA, from 48 hours onwards. Samples were collected every 24 hours for PT gastruloids and every 12 hours for CTCF-AID gastruloids (Figure 2A). Principal component analysis (PCA) showed that gene expression changes associated with gastruloid development explain 47.5% of the variance (Figure 2B, PC1). The treated and untreated gastruloids grown from PT cells were nearly indistinguishable (Figure 2B, in purple), indicating that IAA treatment has a negligible effect on the transcriptome of untagged cells. CTCF-AID gastruloids followed a highly similar trajectory along PC1 whether CTCF was depleted or not (Figure 2B, in orange). These results indicate that cell differentiation is the main driver of gene expression changes in CTCF-AID gastruloids independently of the treatment.

**Figure 2.**
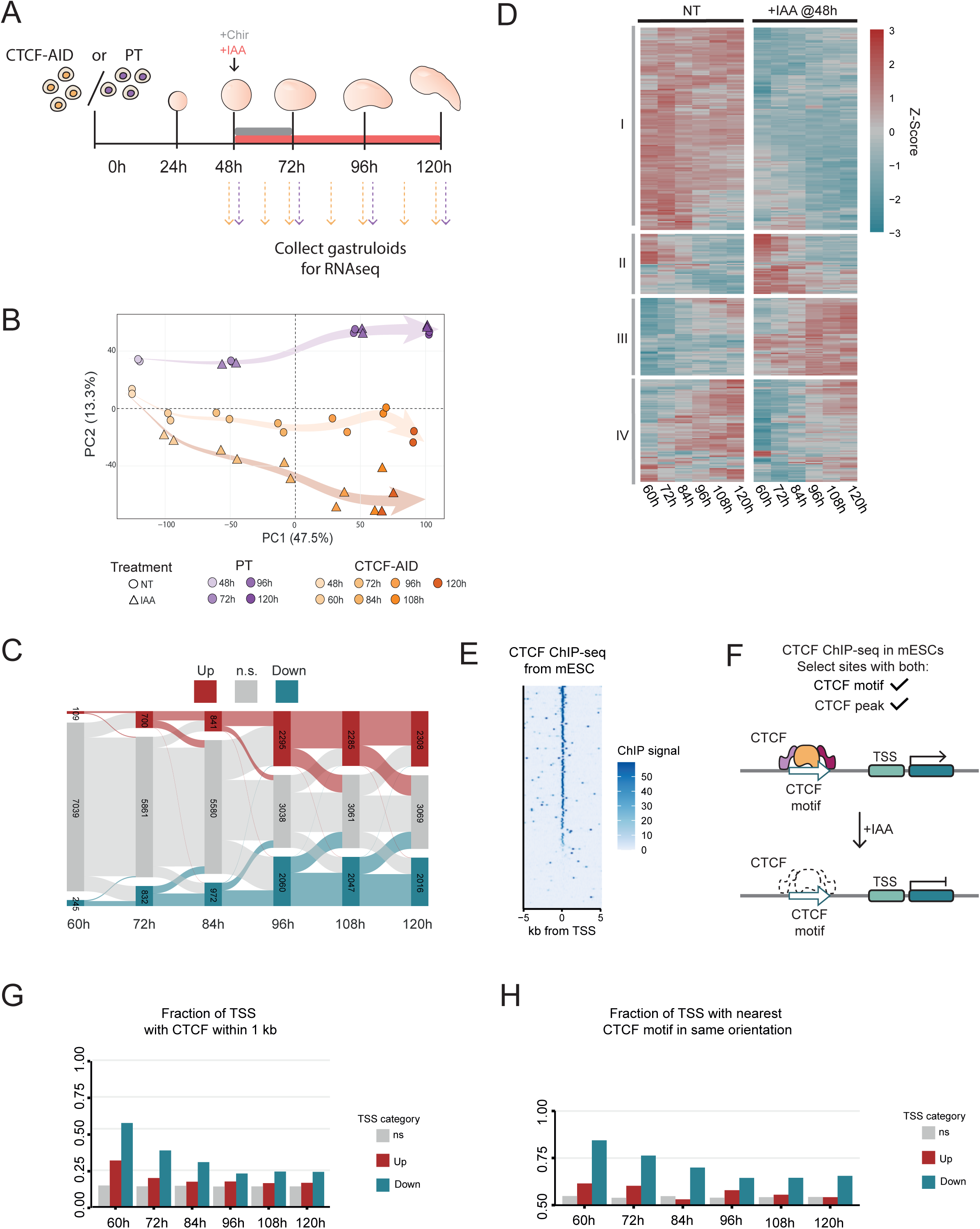
Effect of CTCF depletion on gene expression during gastruloid development A. Schematic representation of the RNA-seq timecourse experiment performed on CTCF-AID and PT gastruloids. +IAA treatment is indicated in red. +Chir stimulation is indicated in grey. Samples were collected every 12h for CTCF-AID gastruloids (orange arrows) and every 24h for PT gastruloids (purple arrows). B. Principal component analysis performed in the RNA-seq samples. PT samples are colored in purple and CTCF-AID are in orange. Circles represent the NT gastruloids and triangles the +IAA@48h treatment. C. Alluvial plot of upregulated (red), downregulated (blue) and non-significant (grey) genes between CTCF-AID NT and CTCF-AID +IAA@48h gastruloids through time. D. Clustered heatmap of differentially expressed genes at 60h and their expression dynamics at later timepoints. E. CTCF Chip-seq signal at the transcription start site (TSS) of downregulated genes at 60h. The Chip-seq data was collected on mESC in Nora et al., 2017^39^. F. Schematic of CTCF sites considered in follow up analysis (G and H). CTCF binding sites needed to have a CTCF motif and a CTCF peak on Chip-seq. G. Fractions of transcription start sites (TSS) of differentially expressed genes (TSS category) at different timepoints of gastruloid development that have CTCF binding within 1kb. H. Fraction of TSS with CTCF binding within 1kb (G) where the CTCF motif has the same orientation as transcription.

We further explored the differentiation trajectories of PT and CTCF-AID gastruloids by deconvoluting the bulk RNA-seq signal using single-cell data from Suppinger et al (2023)^66^ as reference (Figure S2A). This analysis showed that CTCF-depleted gastruloids consisted of the same cell types as the NT and PT gastruloids (Figure S2A), confirming our ATAC-seq observations. However, the deconvolution of the developmental RNA-seq data revealed that CTCF-depleted gastruloids showed subtle differences in the temporal dynamics of the differentiation trajectory. Paraxial mesoderm cells appeared earlier in CTCF depleted gastruloids than in controls, suggesting accelerated development, possibly at the expense of presomitic mesoderm cells, which were not detected at any of the sampled stages.

Increasing the temporal resolution between 60 and 96 hours may clarify the fate of the pre-somitic mesoderm in CTCF depleted gastruloids. Despite the differences in the temporal dynamics, the cell composition at 120 hours of gastruloid development remained similar between control and depleted gastruloids. The only difference at 120 hours was that a primitive streak-like signature persisted in gastruloids lacking CTCF (Figure S2A). Together, these results show that CTCF-depleted gastruloids follow a similar differentiation trajectory as the PT gastruloids. However, our data does not resolve the developmental trajectory of the pre-somitic mesoderm in the absence of CTCF, nor the consequences of altered temporal dynamics for cell types not captured in this model.

Since CTCF-depleted gastruloids failed to elongate, we investigated the expression of key morphogenetic pathways. Notch, Wnt, bone morphogenetic protein (BMP) and retinoic acid (RA) pathways are critical players in the developing embryo and in gastruloids^4,5,13,14,67,68^.

Gene set enrichment analyses showed that CTCF depletion did not affect any of these pathways globally (Figure S2B). We further examined the expression of main drivers of each of these pathways — *T*, *Axin2*, *Wnt3a* and *Sfrp5* for Wnt; *Notch1*, *Nle1*, *Dll1* and *Dll3* for Notch; *Smad5*, *Bmp4*, *Bmp2* and *Nog* for BMP; and *Rara*, *Aldh1a2*, *Cyp26a1* and *Rarb* for RA pathways — to check if their expression was affected either in strength or time of activation (Figure S2C). Gene-specific expression timing and trajectory of these genes remained largely unchanged upon IAA treatment, indicating stable regulatory dynamics. Moreover, the expression level of most genes was broadly comparable in CTCF-depleted gastruloids to the controls. The exceptions were *Wnt3a* and *Nle1*, which had lower expression levels in the CTCF-depleted gastruloids. Overall, the main morphogenetic pathways were properly activated and maintained in the absence of CTCF. Thus, CTCF depletion has a minimal effect on cell differentiation or the temporal activation of morphogenetic pathways during gastruloid development.

### CTCF binds to the promoters of genes downregulated early during gastruloid development

To gain insight into the genes whose expression is controlled by CTCF and could be responsible for the elongation failure, we performed a differential expression analysis along the developmental trajectory comparing CTCF-AID NT vs +IAA@48h gastruloids. We found 109 genes are upregulated and 245 genes are downregulated after 12 hours of IAA treatment (60 hours of gastruloid development) (Figure 2C). At later time points, the number of differentially expressed genes (DEG) gradually increased, reaching 2308 upregulated and 2016 downregulated genes at the 120 hours time point, presumably reflecting the accumulation of secondary effects after prolonged CTCF depletion.

Because CTCF depletion at 48 hours, but not at 72 hours, provokes severe developmental defects, we argued that CTCF-regulated genes involved in gastruloid elongation are likely among the DEGs identified at 60 hours. Therefore, we focused on the DEGs at 60 hours and their expression pattern throughout gastruloid development (Figure 2D). We performed hierarchical clustering and identified four clusters of genes with largely homogeneous intracluster dynamics upon CTCF depletion. Clusters II and IV contain genes whose respective down- or upregulation were delayed upon CTCF depletion. Cluster III is formed by genes that were upregulated upon CTCF depletion, which may be explained by the establishment of ectopic contacts between enhancers and promoters upon CTCF depletion^69^. Finally, cluster I, the biggest cluster containing 162 DEGs at 60 hours, contains genes that were rapidly downregulated following CTCF depletion and remained suppressed throughout the entire gastruloid time course.

To understand how CTCF regulates these globally downregulated genes (cluster I), we examined ChIP-seq data for CTCF^39^ in mESCs. We found that CTCF binds to the transcription start site (TSS) of most of the cluster I genes (Figure 2E). We then systematically quantified CTCF occupancy at the TSS of DEGs by selecting CTCF motifs overlapping a CTCF ChIP-seq peak (Figure 2F). We found that ∼57% of the early downregulated genes at 60 hours contained a bound CTCF motif within 1kb of their TSS (Figure 2G), in contrast to ∼32% of upregulated genes and ∼15% of stable genes. This enrichment gradually decreased over time, likely due to the accumulation of indirectly regulated genes within the downregulated gene sets. Furthermore, we found that within the early downregulated genes with CTCF binding within 1 kb from TSS, ∼84% of them had the nearest CTCF motif in the same orientation as transcription (Figure 2H). In conclusion, CTCF largely promotes gene expression in gastruloids through direct promoter binding.

### CTCF DBD is sufficient for the expression of early downregulated genes

CTCF binds to its DNA motif in an orientation-dependent manner. In fact, TAD boundaries and CTCF-anchored chromatin loops are formed between two convergent CTCF sites bound by CTCF^32,70,71^. This orientation-specific binding allows the N-terminal domain of CTCF to interact with cohesin and results in the unidirectional stalling of loop extrusion^36,40^. Because we observed an orientation bias of the CTCF motif at the TSS of early downregulated genes (Figure 2H), we wondered whether the N-terminal domain is involved in the regulation of these genes by blocking cohesin loop extrusion. The C-terminal domain, on the other hand, has been shown to harbor a negatively charged region that can influence CTCF binding to DNA^72^ and a domain that interacts with RNA^35,73^. To assess the role of the N- and C-terminal domains in gene regulation and gastruloid development we performed reconstitution experiments where we overexpressed CTCF truncations in CTCF-AID cells (Figure 3A). To ectopically express CTCF variants we created an expression plasmid that we randomly integrated into the genome using the PiggyBac transposome system. As positive and negative controls, we integrated the full-length (FL) CTCF or an empty construct expressing only GFP, respectively. The expression of the ectopic factors was controlled by the *EF1a* promoter, which is highly active at all gastruloid stages (Figure S3A). To assess the effect of the exogenous CTCF truncations, we performed bulk RNA-seq on the untreated CTCF-AID mESCs and after depleting the endogenously tagged CTCF for 24 hours. We observed a downregulation of the expected set of genes in the GFP-only expressing cells, whereas cells expressing the FL CTCF rescued the expression of the down-regulated genes (Figure 3B).

**Figure 3.**
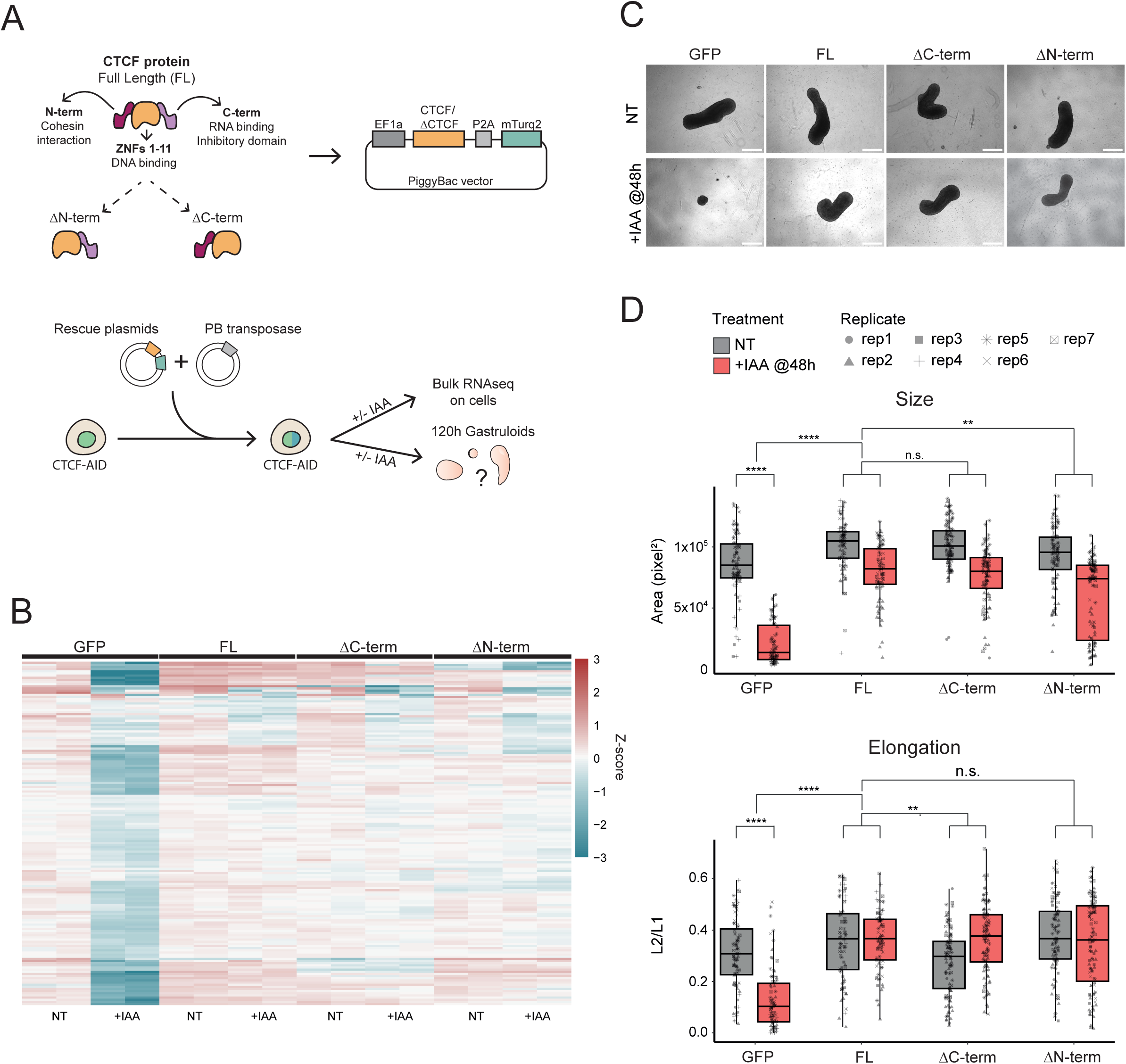
CTCF promoter binding is required and sufficient to rescue gene expression and early gastruloid development. A. Schematic representation of different CTCF protein versions re-introduced on CTCF-AID cells to create the rescue cell lines. The coding sequences were randomly integrated in the genome under the EF1a promoter. mTurquoise2 (mTurq2) was used as a selection marker and to control for the expression levels of the re-expressed proteins. B. Heatmap showing the Z-score values of the genes downregulated at 60 hours upon CTCF depletion (same set of genes as in Fig2B, cluster I) for each of the reconstituted cell lines. GFP is a negative control where GFP was introduced in the cells instead of a CTCF version. Upon treatment, the genes are downregulated in the GFP-only line but their expression was rescued in the reconstituted cell lines. Data was collected on mouse embryonic stem cells. +IAA indicates treatment with auxin for 24 hours. C. Representative brightfield images of 120 hours gastruloids from the GFP-only and reconstituted cell lines. D. Quantification of morphological features on the brightfield images of 120 hours gastruloids. Size was measured by calculating the area on pixel^2^. Elongation was measured as the ratio between the L2 and L1 LOCO-EFA coefficients^95^. Values were obtained using MOrgAna^94^. Difference in treatment effects was tested using linear models with Benjamini-Hochberg multiple testing correction (n.s. = p > 0.05; * = p ≤ 0.05; ** = p ≤ 0.01; *** = p ≤ 0.001; **** = p ≤ 0.0001).

To our surprise, both the ΔN-terminal and the ΔC-terminal CTCF variants rescued the expression of the majority of the downregulated genes. These results indicate that the CTCF DNA binding domain (DBD) is required and sufficient to drive the expression of these genes. Furthermore, the observation that the ΔN-terminal CTCF mutant rescues the expression of genes bound by CTCF at the promoter is inconsistent with a role of cohesin-mediated loop extrusion in the regulation of these genes.

### CTCF DBD and N-terminal domain have different roles in gastruloid development

Having established a cell line system in which we can switch from the endogenous CTCF to a truncated variant, we proceeded to examine the effect on gastruloid development. We performed the gastruloid protocol and found that in the untreated condition we were able to generate gastruloids from every cell line (Figure 3C, NT). Given that the ΔN- and ΔC-terminal mutants rescued CTCF promoter regulation, we wondered whether these CTCF truncations were also sufficient to rescue gastruloid development. To this end, the variant-expressing gastruloids were treated with IAA from 48 hours (Figure 3C). As expected, the GFP-only expressing gastruloids failed to elongate (Figure 3C-D) and were significantly smaller than their untreated counterpart (Figure 3C-D). Adding back the FL CTCF yielded elongated gastruloids with a similar size as the NT controls (Figure 3C-D). Furthermore, both ΔN- and ΔC-terminal mutants rescued gastruloid development (Figure 3C-D). These gastruloids were elongated and bigger than the GFP-only treated gastruloids (Figure 3C-D). Taken together, these findings indicate that the CTCF DNA-binding domain is sufficient for rescuing gastruloid elongation and the maintenance of the gene regulatory network necessary for early gastruloid morphogenesis.

While we show that CTCF promoter binding is crucial for early *in vitro* development, the role of CTCF as a regulator of chromatin looping has been shown to be crucial for embryonic development^23,26,74^. To determine whether the extrusion stalling function of CTCF plays a role in later gastruloid development, we performed experiments where we prolonged gastruloid culture until 168 hours (Figure 4A). Consistent with the results at 120 hours (Figure 3), GFP-only gastruloids treated from 48 hours failed to develop, while the full-length CTCF reconstituted gastruloids were elongated albeit smaller under the same treatment conditions (Figure 4B-C). Reintroducing the ΔC-terminal CTCF truncation largely rescued the elongation with a partial rescue in size (Figure 4B-C). Interestingly, gastruloids expressing the ΔN-terminal CTCF truncation collapsed between 120 hours and 168 hours (Figure 4B). Their size was drastically reduced and instead of creating a polarized elongated structure, they presented an oval shape (Figure 4C). These results clearly show that while the extrusion stalling function of CTCF is not required in the early stages of gastruloid development, it becomes crucial at later stages.

**Figure 4.**
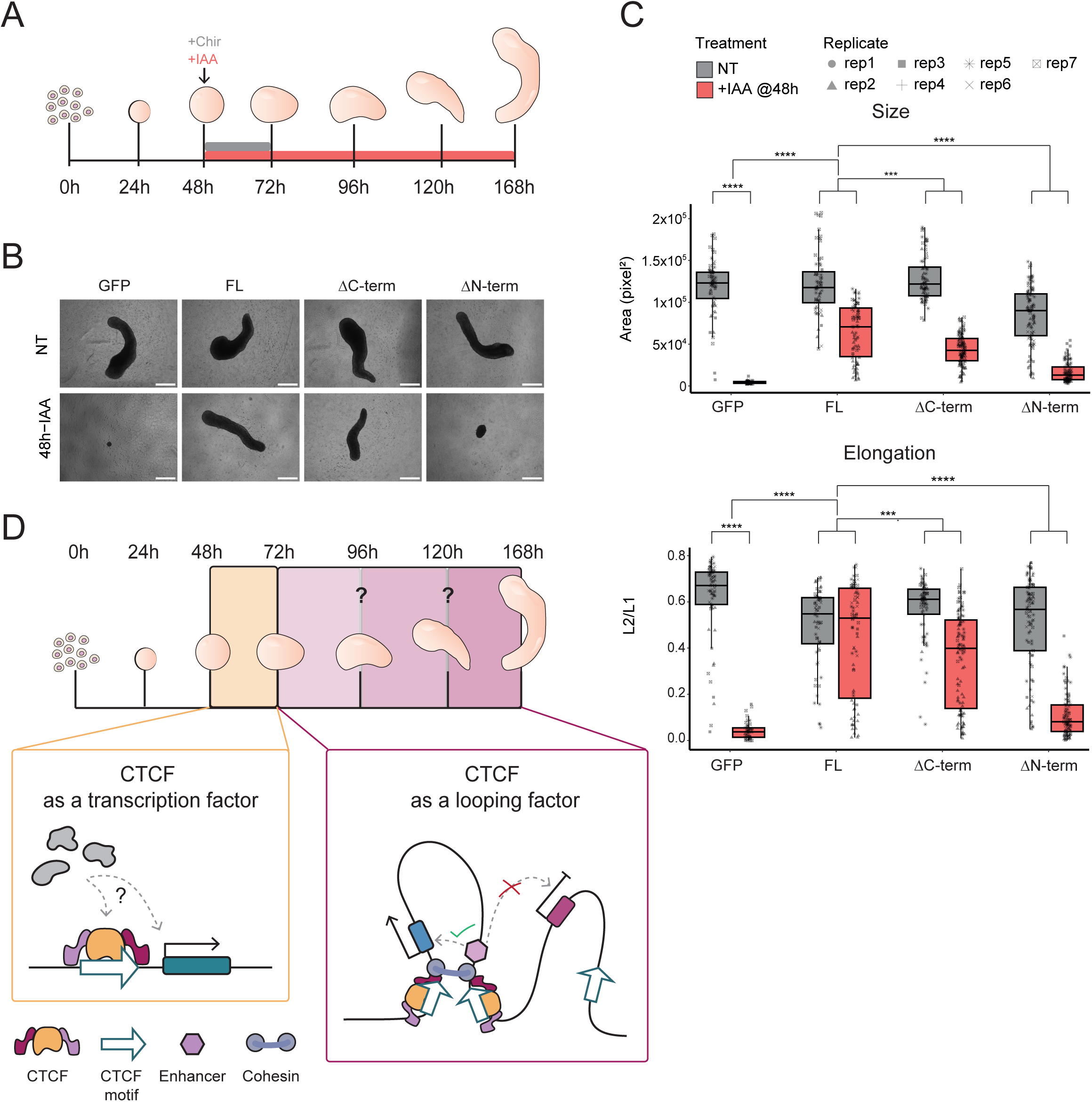
CTCF mediated extrusion stalling is required for late gastruloid development. A. Schematic representation of prolonged gastruloid culture and treatment regime. B. Representative brightfield images of 168 hours gastruloids for the GFP and reconstituted cell lines. C. Quantification of morphological features on the brightfield images of 168h gastruloids. Size was measured by calculating the area on pixel^2^. Elongation was measured as the ratio between the L2 and L1 LOCO-EFA coefficients^95^. Values were obtained using MOrgAna^94^. Difference in treatment effects was tested using linear models with Benjamini-Hochberg multiple testing correction (n.s. = p > 0.05; * = p ≤ 0.05; ** = p ≤ 0.01; *** = p ≤ 0.001; **** = p ≤ 0.0001). D. Model of the dual function of CTCF in morphogenesis. Between 48h and 72h of gastruloid development CTCF needs to bind at the promoter of target genes to promote their expression and gastruloid development. At later stages CTCF looping activity is essential to maintain the proper gene regulatory network during morphogenesis and avoid ectopic gene expression.

## Discussion

Using an *in vitro* model of early mouse embryonic development combined with rapid depletion of CTCF, we were able to uncouple morphogenesis from cellular differentiation. Our results show that two discrete functions of CTCF contribute to gastruloid morphogenesis at distinct phases in development (Figure 4D). The morphogenetic defects observed upon CTCF depletion are likely driven by a subset of genes that become downregulated in the absence of CTCF in the early phases of the gastruloid time course. The CTCF DNA-binding domain is both essential and sufficient to sustain the expression of these genes, likely through direct binding at their promoter regions. The looping function of CTCF is required during the later stages of gastruloid development.

While in regular 2D monolayer culture CTCF depletion shows limited direct effects on gene expression^31,34,39,58^, in gastruloids CTCF is essential for proper morphogenesis. This highlights the usefulness of 3D *in vitro* models where morphological effects can be used as a readout. Our experiments conclusively showed that promoter binding by CTCF is critical for the expression of a relatively small number of genes, which play an important role in gastruloid development. These genes are enriched for CTCF binding at the promoter. The CTCF motif at these promoters occurs in the same orientation as that of transcription.

Similarly to our CTCF-depleted gastruloids, mouse embryos deficient for CTCF fail to gastrulate^57^. They also present CTCF binding at the TSS of downregulated genes^57^, which may explain the lethality observed in these embryos. Importantly, in human embryos^29^, loss of CTCF also leads to downregulation of promoter-bound genes, implying that promoter-associated CTCF activity may likewise be required for human embryonic development. Together, these observations point to a conserved role for CTCF-mediated transcriptional regulation at promoters in mice and humans. Upon inspection of the target genes, there were no obvious candidates to explain its role specifically in gastrulation. An open question remains which CTCF target gene is the critical gene for gastruloid formation, or whether multiple target genes are required to drive this process. Furthermore, we showed that the DNA binding domain of CTCF alone is enough to rescue the expression of promoter-bound genes, but it is unclear how the zinc finger array may result in the activation of genes. Molecular and biochemical experiments are necessary to uncover how CTCF promotes the expression of these genes. This study suggests a cohesin independent function of CTCF regulating promoter-bound promoters, in agreement with previously published work^39,75^.

Despite its critical role in gene activation and gastruloid formation, CTCF is not required for cell fate specification during gastruloid development. These observations are substantiated by experiments in other model systems. Mouse embryos lacking CTCF can go through the first wave of differentiation and progress until the morula stage^57^. Moreover, transdifferentiation of human B cells into macrophages can also progress in the absence of CTCF^58^. While CTCF is required for extrusion stalling we have recently shown that loss of the extrusion factor NIPBL does not impair cellular differentiation either in 2D monolayer culture nor in gastruloids^76^. These findings suggest that large-scale genome organization and loop extrusion are not strictly required for cell state transitions.

Our CTCF reconstitution experiments revealed that the N-terminal domain of CTCF plays an important role in post-gastrulation development. The N-terminal domain of CTCF plays a crucial role in the formation of loops and TADs, which are essential for proper organismal development. For instance, CTCF-dependent boundaries are crucial for proper brain^50^ and limb^51^ development. Furthermore, the temporal and spatial activation of the *Hox* genes in gastruloids is regulated by CTCF boundaries^26^. The looping function of CTCF is mediated through the interaction of a YDF amino acid sequence in the N-terminal domain with cohesin^36,40^. Interestingly, the homozygous mutation of this peptide sequence is embryonically lethal in mice, underscoring the importance of the extrusion stalling function in development *in vivo*^77^. In 2D monolayer culture, however, mutations in the YDF domain have limited consequences on cellular fitness^36,77^. Our experiments indicate that the phenotypic consequences of extrusion stalling occur post-gastrulation. However, further experiments are necessary to pinpoint the timepoint at which the consequences of a defect in extrusion stalling become apparent, which may in fact lie before gastrulation.

The emergence of stem-cell based embryo models, like gastruloids, is transforming the study of embryonic development^4–8,10,11,13–15,26,67,78,79^, by providing an accessible framework to probe the functional role of specific factors in a controlled multicellular developmental context. In the current study we have used them to uncover two complementary functions of CTCF in morphogenesis. While the organization of the genome into CTCF-anchored chromatin loops is unlikely to be important for lineage specification, it is critical for the fine-tuning of expression of key developmental factors. Although perturbations of CTCF-anchored loops may not have catastrophic effects on early preimplantation development, they likely influence phenotypic variation in ways that could affect individual fitness. Gastruloids present a scalable model system to investigate the role of the CTCF insulation and loops in development to better understand its function in early development. The recent development of more advanced *in vitro* embryo models^80–82^ will likely enable us to probe the role of CTCF and looping in early development with even more precision.

## Resource Availability

### Lead contact

Further information and requests for resources should be directed to and will be provided by the lead contact, Elzo de Wit.

### Materials availability

Plasmids and cell lines generated in this study are available upon request.

### Data and Code availability

- ATACseq and RNA-sed data have been deposited to Gene Expression Omnibus (GEO) database: GSE313533.
- Imaging data, Json files containing the raw morphological quantifications, GSEA results and FCS files will be available on Zenodo as of the date of publication (DOI: 10.5281/zenodo.17911392).
- This paper does not report original code.
- Any additional information required to reanalyze the data reported in this paper is available from the lead contact upon request.

## Acknowledgements

We thank the NKI Flow Cytometry Facility, NKI Genomics Core Facility, NKI Bioimaging Facility and NKI Research High Performance Computing Facility. We thank Teun van de Brand for initial ATACseq analyses. We thank members of the de Wit lab for critically reading the manuscript. Work in the de Wit laboratory is supported by the Dutch Research Council (016.161.316, Vidi; VI.C.222.049, Vici) and the European Research Council (637587, ‘HAP-PHEN’; 865459, ‘FuncDis3D’). Research at the Netherlands Cancer Institute is supported by an institutional grant of the Dutch Cancer Society and of the Dutch Ministry of Health, Welfare and Sport.

## Author Contributions

N.A.S., M.M., L.B. and E.d.W. conceived and designed the study. N.A.S., L.B. and M.R. performed the experiments. N.A.S. and M.M. analysed data. E.d.W. and L.B. supervised the study. P.S generated microwells under the supervision of S.G.. N.A.S., M.M., L.B., and E.d.W. wrote the manuscript with input from all authors. All authors read and approved the manuscript.

## Methods

### Key Resource table

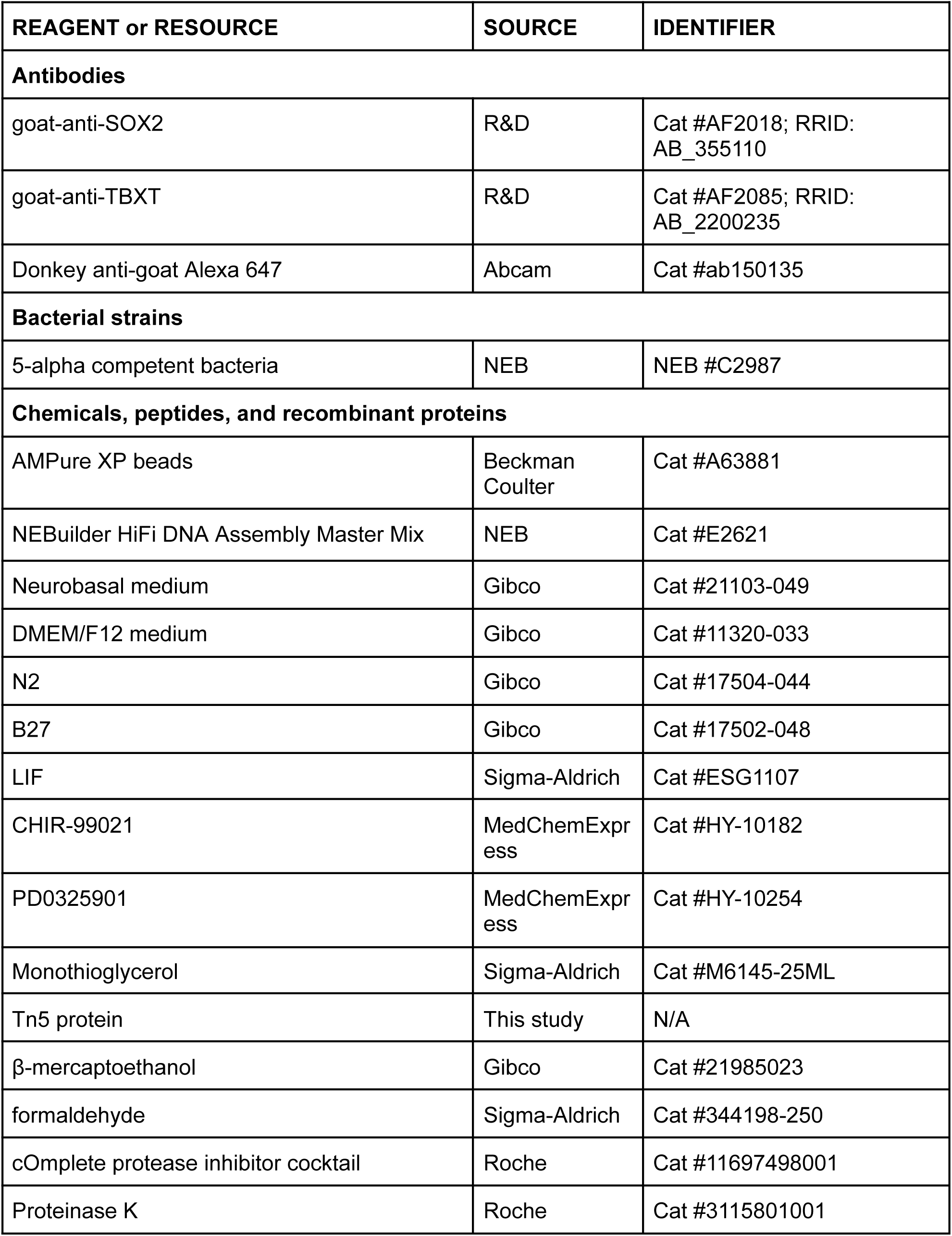
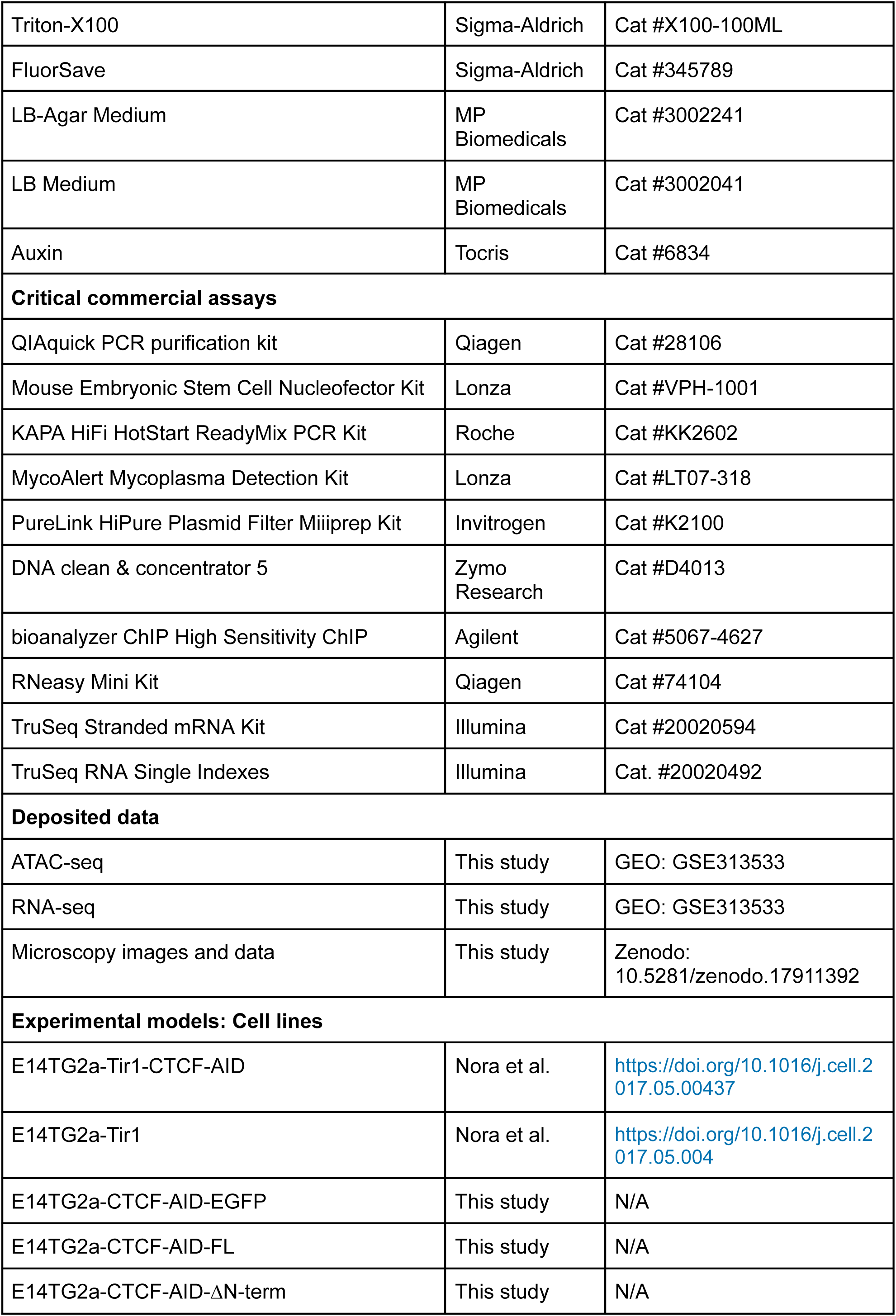
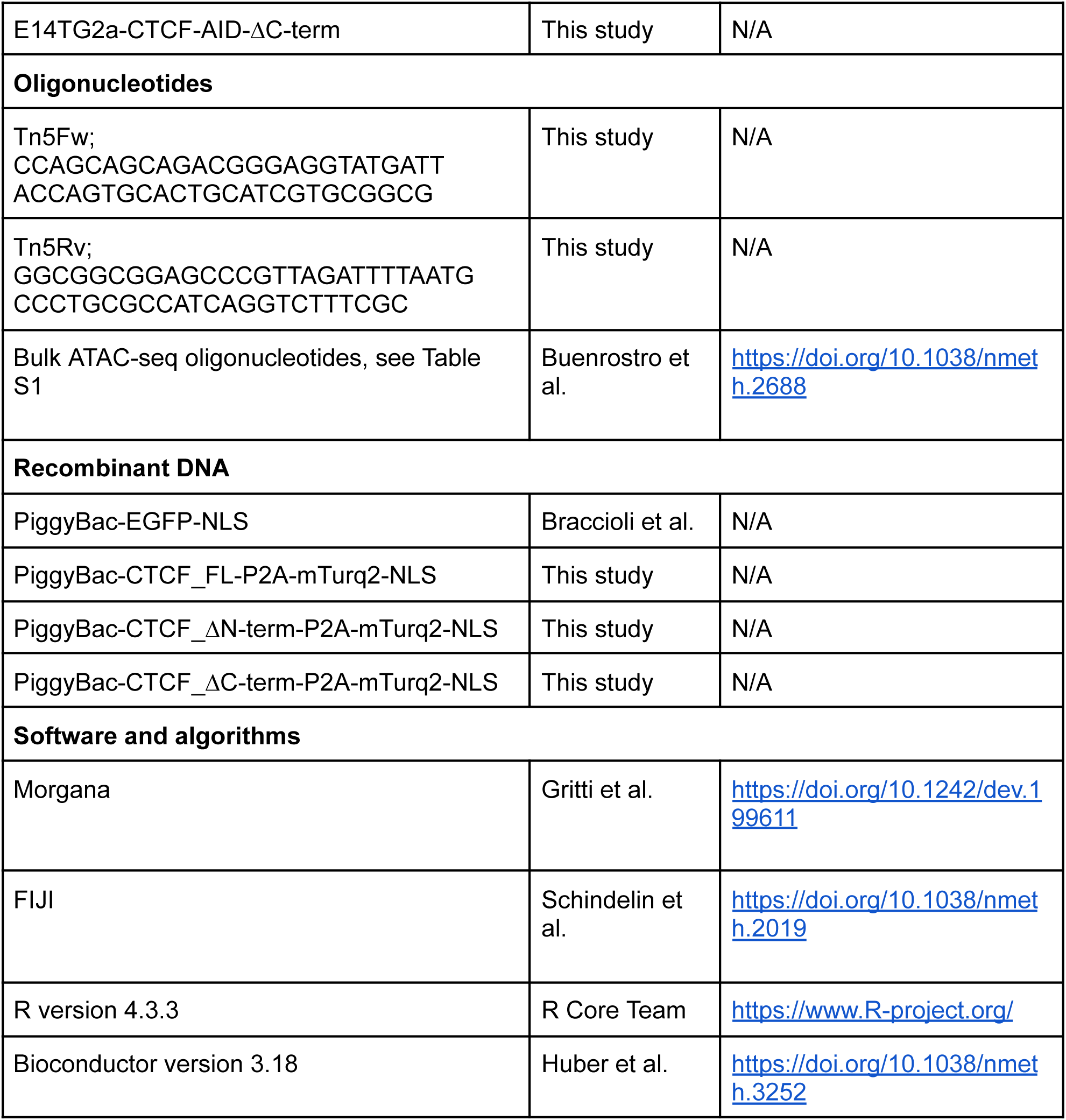

### Experimental model and study participant details

#### Cell lines

CTCF-AID mESCs (E14TG2a) and PT mESCs were obtained from Elphege Nora’s lab. The CTCF-AID cells were utilized to create the CTCF Full Length, CTCFΔN-terminal and the CTCF ΔC-terminal reconstituted cell lines. All mESCs were cultured in 2i-LIF medium with 50% DMEM/F12 (Gibco) and 50% Neurobasal media (Gibco) supplemented with 0.5x N2 (Gibco), 0.5x B27 + retinoic acid (Gibco), 0.05% BSA (Gibco), 3μM CHIR99021 (BioConnect), 1μM PD03259010 (BioConnect), 2mΜ Glutamine (Gibco), 1.5×10-4 M 1-thioglycerol (Sigma-Aldrich), 100U/ml LIF (Cell guidance systems) and 50U/ml penicillin-streptomycin (Gibco) on gelatin-coated 10cm petri dishes in a humidified incubator (5% CO2, 37°C).

To generate the reconstituted cell lines, CTCF-AID cells were electroporated with 1.8ug of PiggyBac-transposase vector (mPB-L3-ERT2-mCherry) and 18ug of PiggyBac vectors (gifted by Bas van Steensel) containing different cargos: EF1a promoter with EGFP fused with a nuclear localization signal (NLS) or EF1a promoter with different versions of the CTCF coding sequence followed by a self cleaving peptide (P2A) and mTurquoise2 fused with a NLS. The day after electroporation, cells were stimulated with 4-OHT for 24h. Cells were expanded and a pool of fluorescently labeled cells was sorted by flow cytometry.

Mouse Embryonic Stem Cell Nucleofector Kit (Lonza) was used for electroporation of mESCs on an Amaxa Nucleofector II device (Lonza) according to manufacturer instructions. The cell pools were used for gastruloid culture and sequencing experiments.

#### Gastruloid culture

Gastruloid culture was performed as previously described^5,13^. Cells were dissociated in differentiation medium using a 5ml serological pipette attached with a 200μl pipette tip. The differentiation medium consists of DMEM/F12 and Neurobasal medium supplemented with 0.5X N2, 0.5X B27 + Retinoic acid, 2mΜ Glutamine, β-mercaptoethanol and 50U/ml penicillin-streptomycin. Approximately 390 single cells were planted per well in U-bottom 96-well plates (Thermo Scientific) with 40μl of differentiation medium and let aggregate for 48h in a humidified incubator (5% CO2, 37°C). Next, gastruloids were stimulated for 24h with 3μM CHIR99021 in 150μl of differentiation medium using a Hamilton Star R&D liquid handling platform attached to an incubator. The medium was manually refreshed every 24h with 150μl of differentiation medium. Gastruloids were cultured until 120h or until 168h. For the extended gastruloid culture until 168h, gastruloids were kept in a classical humidified incubator (5% CO2, 37°C) and the differentiation medium was changed every 24h. Pictures were taken using an inverted wide field microscope (Zeiss Axio Observer Z1 Live) at 120h and 168h. For immunostaining, gastruloids were fixed with 4% formaldehyde (Sigma-Aldrich) at approximately 60h and 72h, washed with PBS once and stored in PBS at 4°C. For bulk ATAC sequencing, gastruloids were collected at 120h. For RNA-seq, gastruloids were cultured in elongated thin-walled microcavities with U-shaped bottoms^83^. PT gastruloids were collected every 24h while CTCF-AID gastruloids were collected every 12h in the RLT (RNA purification kit, Qiagen) buffer from and including 48h to 120h. In all sequencing experiments gastruloids were pooled per condition, dissociated into single cells and stored at −80C. Gastruloid dissociation was performed by mild trypsinization and gentle resuspension with a 200μl tip.

### Method details

#### Cloning

All plasmids used in this study were created by gibson assembly^84^. The full CTCF coding sequence (1-735 amino acids) was amplified by PCR using pKS004-pCAGGS-3XFLAG-CTCF-eGFPk^40^ was a gift from Elphege Nora (addgene plasmid #156438) as template. The N-terminal (1-265 amino acids) deletion CTCF coding sequence + P2A was synthesized by Twist Bioscience. The fragment was further amplified by PCR with primers compatible for gibson ligation. The C-terminal (576-735 amino acids) deletion CTCF sequence was obtained by PCR amplification using the full coding sequence as template. The mTurquoise2 sequence was PCR amplified from pME034^85^ (a gift from Van Steensel) with gibson primers. The PiggyBac backbone vector containing the EF1a promoter and the NLS sequence was linearized by PCR from PiggyBac-EGFP-NLS^5^. The fragments were ligated together with the vector in a one step ligation reaction with the NEBuilder HiFi DNA Assembly Master Mix (NEB #E2621) following the manufacturer guidelines. 5*α* competent bacteria (NEB #C2987) were transformed with the ligation product and plated on 10cm plates coated with LB agar with ampicillin (0.5 μg/ml) for antibiotic selection. Picked colonies were cultured in suspension LB overnight supplemented with ampicillin. DNA was isolated with the PureLink HiPure Plasmid Filter Midiprep Kit (Invitrogen, K2100). Plasmids sequences were checked by whole plasmid sequencing (Oxford Nanopore by Macrogen).

#### Bulk ATAC-seq

The transposome was assembled as previously described^5^. Briefly, the transposons were annealed by mixing 10μl of 10X TE buffer with 45μl of ME-REV (100μM in H2O) and with either 45μl of ME-A (100μM in H2O) or ME-B (100μM in H2O) (see Table S1 for details). Next the adapter solution was incubated at 95°C for 10min and cooled down to 4°C at 0.1°C/sec. For transposome formation, the annealed adapters were diluted in equal volume of H2O. The Tn5 protein was purified by the NKI protein facility as previously described^5^. The adapters were incubated with 0.2mg/mL Tn5 in a 1:20 ratio for 1h at 37°C. The annealed transposome was used directly or stored at −20°C for no longer than 2 weeks.

Previously frozen gastruloid cell suspensions were rapidly thawed and 50,000 cells were resuspended in cold PBS and lysed with a 2x lysis buffer (Tris-HCl pH7.5 1M, NaCl 5M, MgCl2 1M, 10% IGEPAL). Then, cells were centrifuged and the pellet was incubated with 2 μl of transposon mix (A and B in equal ratios) and 2xTD buffer (20 mM Tris(hydroxymethyl)aminomethane; 10 mM MgCl2, 20% dimethylformamide, brought at pH 7.6 with acetic acid). Tagmentation was performed at 37°C for 1h. Subsequently, samples were incubated at 40°C for 30min with a clean-up buffer (55% buffer 1 (5M NaCl, 0.5M EDTA), 5% SDS and 10mg/ml ProtK). The sample was further purified by fragment selection with 2X AMPure XP beads. The supernatant was amplified by two consequent rounds of PCR using KAPA HiFi HotStart ReadyMix with P5 and P7 indexed primers (see oligonucleotide list). Fragments between 200 and 700 bp were purified using AMPure XP beads. Quality control was performed by Bioanalyzer High Sensitivity DNA analysis (Agilent).

#### RNA-seq

Cell suspensions from the gastruloid time course stored in the RLT buffer at −80°C were used for RNA purification with the Qiagen RNeasy Mini kit (Qiagen), including a DNaseI treatment. The purified RNA was used for library preparation using the TruSeq Stranded mRNA kit (Illumina) with TruSeq RNA Single Indexes set A and B (Illumina) and following the standard protocol.

#### IF and image processing

The previously fixed gastruloids were immunostained as described previously^5^. Briefly, gastruloids were blocked with PBS-FT (10% fetal bovine serum and 0.2% Trixton-X100 in PBS) for 1h at room temperature. Overnight incubation at 4°C with the primary antibodies was performed in the PBS-FT buffer with antibodies against SOX2 (1:50; AF2018) or TBXT (1:250; AF2085). Next, gastruloids were washed three times with PBS-FT and incubated with the secondary antibody Alexa 647 (1:250; Ab150135) overnight at 4°C. Finally, the secondary antibody was washed three times and the gastruloids were mounted in microscope slides using FluorSave (Sigma Aldrich). Gastruloids were imaged in a SP8 confocal microscope (Leica). A z-stack was collected for each gastruloid. FIJI^86^ was used for creating a max projection and displaying pixel intensity-based Look-Up-Tables.

### Analysis

#### ATAC-seq data processing and analysis

ATAC-seq data were mapped to the mm10 genome using bwa-mem^87^ (v0.7.17-r1188), and samtools^88^ (v1.19.2) was used to filter properly paired and uniquely mapped reads with MAPQ>10, and to perform duplicate read removal. To deconvolve the contributions of constituent cell types in bulk samples, a reference gastruloid single-cell ATAC-seq dataset was used^5^. Independent component analysis (ICA) was performed on normalized pseudobulk reference counts as described in Magnitov et al 2025^65^, using the FastICA algorithm with 7 components, implemented in the ica R package (https://cran.r-project.org/web/packages/ica/index.html). Read counts of the query bulk samples in the reference peak set were computed and the unmixing matrix obtained from the ICA was used to unmix the cell type contributions. Finally, the contributions were scaled to the sum of the contributions for each sample to obtain cell type proportions.

#### RNA-seq data processing and analysis

RNA-seq data were mapped to the mm10 genome using STAR^89^ (v2.7.9), and samtools^88^ (v1.19.2) was used to filter properly paired and uniquely mapped reads with MAPQ>10. Read counts in genes were generated using the summarizeOverlaps function from the R/Bioconductor package GenomicAlignments^90^ with the counting mode argument set to “IntersectionStrict”. Read count normalization and differential expression testing at each timepoint was performed using DESeq2^91^ with Wald test, and principal components were computed on the VST-transformed count matrix.

To deconvolve cell type contributions in bulk RNA-seq samples, a reference scRNA-seq dataset was used^66^. Provided cell type annotations were used to aggregate the single-cell counts into a pseudobulk reference count matrix. The resulting matrix was merged with the bulk counts and transformed using VST from DESeq2. Cell type contributions within each bulk sample were decomposed using non-negative least squares, implemented in the nnls R package (https://cran.r-project.org/web/packages/nnls/index.html), on the VST pseudobulk matrix. Final cell type proportions were obtained by scaling to the sum of the contributions for each sample.

#### CTCF binding analysis

Previously published CTCF ChIP-seq data in CTCF-AID mESCs^39^ were re-mapped to the mm10 genome using bwa-mem^87^ (v0.7.17-r1188), and samtools^88^ (v1.19.2) was used to filter properly paired and uniquely mapped reads with MAPQ>10. CTCF signal peaks were called using macs2^92^ with default parameters. CTCF motifs and their orientations in the mm10 genome were obtained from the JASPAR 2024 database^93^. CTCF binding sites were defined by overlapping CTCF motifs with CTCF ChIP-seq peaks using GenomicRanges^90^, and these sites were further used to quantify CTCF binding around gene TSSs.

#### GSEA

The fgsea package in R was used to perform gene set enrichment analysis with *Mus musculus* gene sets from MSigDB. For each gene and timepoint, differential expression analysis yielded log₂ fold-change estimates, which were used as the ranking statistic for enrichment. Gene sets from the reactome canonical pathways collection were analyzed. Fgsea was run with 10,000 permutations, a minimum gene set size of 15 and a maximum size of 1,500. P-values were adjusted for multiple testing using the Benjamini-Hochberg procedure within each timepoint.

#### Morphological analysis

Gastruloid bright field images were masked and analyzed using MOrgAna^94^. The model for image masking and binarization was trained with ∼5% of the images as recommended by the developers. Each of the masks were individually inspected for accuracy and if needed, masks were adjusted manually. The morphological parameters were calculated on the unprocessed mask. To describe the complex shapes of gastruloids we used Lobe-Contribution Elliptic Fourier Analysis (LOCO-EFA) as previously described^95^. Briefly, LOCO-EFA decomposes a 2D shape outline into a series of modes. Each mode corresponds to an “n-lobe” component of the shape. The coefficient Ln quantifies the contribution of the mode with n lobes to the shape. L1 and L2 coefficients were used to describe the shape of gastruloids. L1, is a measure of the overall linear size of the shape while L2 reflects bi-lobed deviations (elongation). To remove the confounding effect of the gastruloid size on the shape, we divided the L2 coefficients by their respective L1. Thus equally elongated gastruloids with different sizes have similar L2/L1 ratios.

Statistical analyses were performed in R on the log-transformed area measurements and L2/L1 LOCO-EFA coefficient ratios. To assess whether the auxin treatment effect at 48h or 72h of gastruloid development differed between the CTCF-AID and parental lines, linear models including an interaction term between treatment and line were fitted. For the CTCF variant re-expression experiments, within each time point the main auxin treatment effect was first tested in the GFP control to confirm the expected baseline phenotype of CTCF depletion. To compare treatment effects across CTCF variants, linear models including an interaction term between variant and treatment were fitted. The interaction terms were tested to assess whether GFP, ΔN-terminal, ΔC-terminal re-expression altered the treatment response relative to full-length CTCF re-expression. P-values were corrected for multiple testing using the Benjamini-Hochberg procedure

---

**Figure S1.**
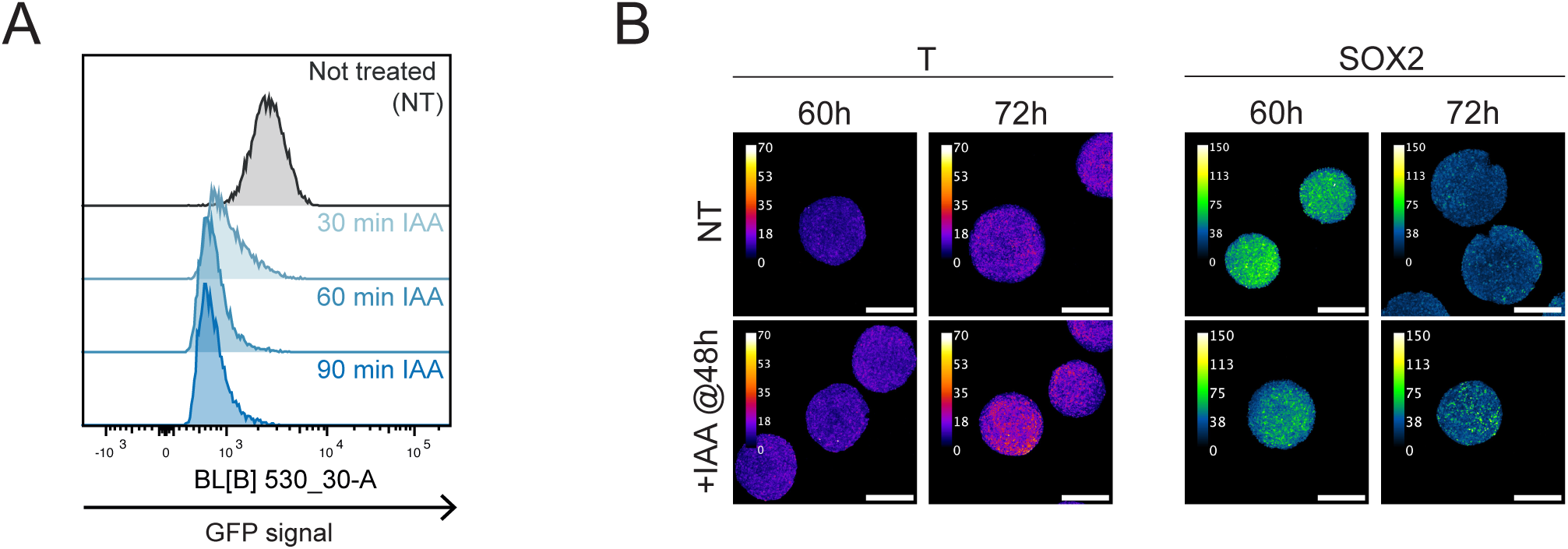
CTCF depletion in mESC and symmetry breaking events in CTCF-AID gastruloids A. GFP signal on CTCF-AID cells measured by fluorescence activated cell sorting during a treatment time course. B. Brachyury (T) and SOX2 immunofluorescence stainings on 60h and 72h gastruloids. Scale bar: 300um. Calibration bar: pixel intensity values on max-projection.

**Figure S2.**
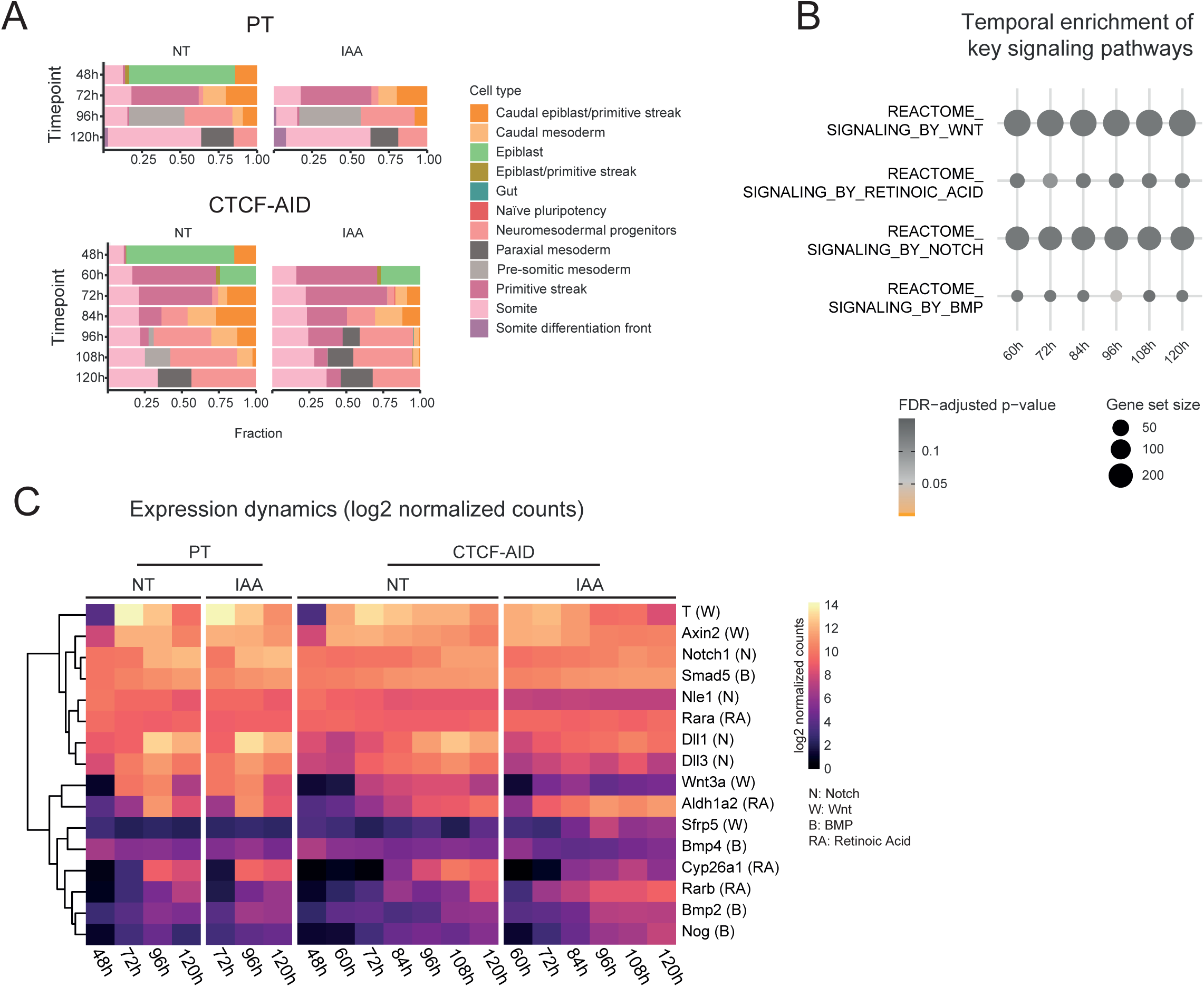
CTCF depletion does not affect the gene expression dynamics of morphogenetic pathways in gastruloids A. Bulk RNA-seq timecourse deconvolution (data from Fig2). Single cell data was published in Suppinger et al 2023^66^. B. Pathway enrichment analysis of key morphogenetic pathways in gastruloids. Gene sets were extracted from Reactome. C. Clustered heatmap showing the log2 normalized counts for genes on the Notch (N), Wnt (W), BMP (B) and retinoic acid (RA) pathways. Clustering was done based on expression values. Data has been grouped based on the cell line (PT and CTCF-AID) and the treatment (Not treated - NT and +IAA@48h).

**Figure S3.**
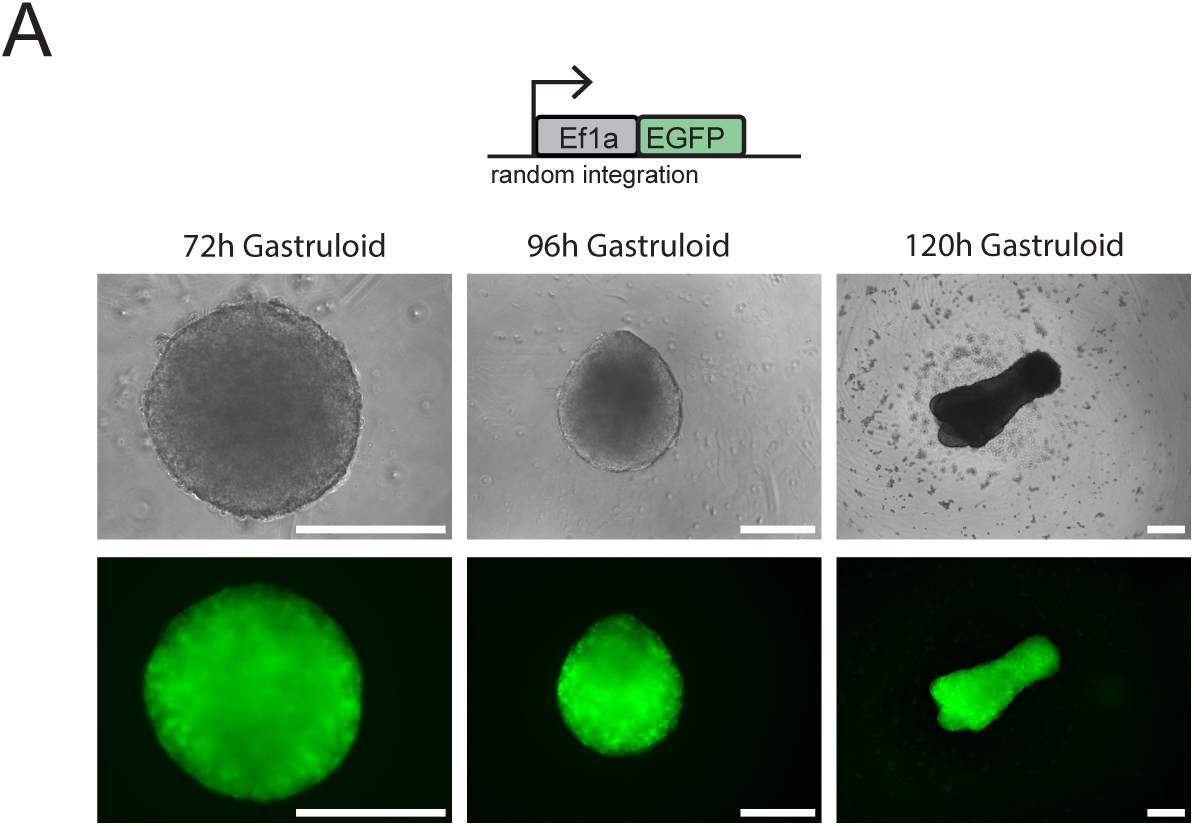
EF1a promoter is active through all gastruloid developmental stages. A. Representative brightfield and fluorescence images (green) of gastruloids at different stages of development. EGFP has been randomly integrated in the genome of the CTCF-AID cells and its expression is driven by the EF1a promoter. Scale bar: 300um.

**Supplementary Table 1.**
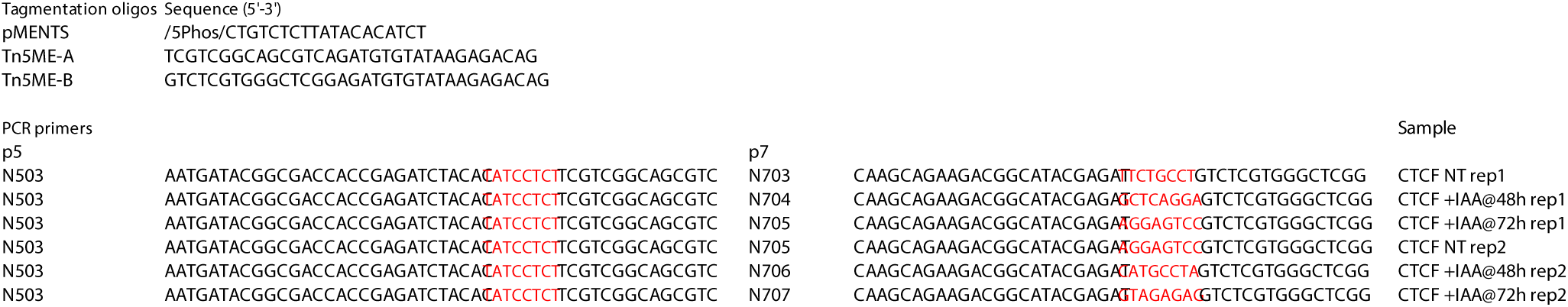
Bulk ATAC-seq oligos.

